# Depletion of a neural cell adhesion molecule disrupts both epithelial and germ layer identity in sea anemone embryos

**DOI:** 10.1101/2025.08.03.668367

**Authors:** Anna Postnikova, Julissa Tello, Maristhela Alvarez, Grace Lee, Ilhan Ali, Christine M Field, Timothy J Mitchison, Katerina Ragkousi

## Abstract

Assembly of cells into epithelial layers marks the early steps of tissue organization and embryonic development in most animals. Although several conserved proteins are known to be essential for epithelialization in bilaterians, it is unclear whether these are sufficient to drive epithelial organization in all metazoans. Using proteomics and knockdown approaches in embryos of the early-branching sea anemone *Nematostella vectensis*, we identified the neural cell adhesion molecule NCAM2 to be essential for organization of the primary epithelial layer. NCAM2 belongs to the immunoglobulin family of cell adhesion molecules with roles in inter-cellular adhesion and signaling. In this work, we show that NCAM2 is enriched at the apical cell junctions and is required for *Nematostella* epithelial cell identity. Importantly, embryos depleted for NCAM2 fail to gastrulate. Both whole-embryo germ layer patterning and tissue changes required for gastrulation are impaired. Together, our data show that in *Nematostella* NCAM2 is required for both epithelial and germ layer organization.

## Introduction

The formation of an epithelial cell layer is among the earliest events in the development of most animals (1, 2). The primary epithelium provides the developing embryo with a permeability barrier that segregates its internal structures from the external environment. One of the most critical subsequent events is gastrulation, during which cells move into the interior of the embryo. Concurrent with gastrulation, the single-cell layered blastula transitions into the gastrula, the stage characterized by the formation of separate germ layers. These layers have distinct developmental trajectories, with the ectoderm in the exterior and the mesoderm and endoderm comprising the interior of the embryo (3, 4).

The sea anemone, *Nematostella vectensis*, is an interesting system for studying the properties of the primary epithelium. As a representative cnidarian, *Nematostella* provides insights into the common ancestor of metazoans (5–7). Furthermore, the simple morphology of the early *Nematostella* embryo draws attention on how epithelial organization determines embryo organization. *Nematostella* embryos also provide an extreme example of cell cycle regulation of epithelial organization, where apical cell polarity is disrupted during mitosis and re-established during interphase (8, 9). Given this background, it is particularly important to examine how the primary epithelium in *Nematostella* embryos is organized at the molecular and cellular level.

Cell-cell adhesion molecules, especially cadherins, play a central role in epithelial organization (10–12). These widely conserved proteins are present in *Nematostella*, but it is unclear whether additional cell-cell or cell-matrix adhesion proteins are present and important (13, 14). Here, we report an unbiased approach to identify surface proteins expressed in the primary epithelia of *Nematostella* embryos. Surprisingly, we found an abundance of proteins resembling neural cell adhesion molecules (NCAMs) (15). These proteins belong to the superfamily of immunoglobulin cell adhesion molecules (IgCAM). Various IgCAM orthologues have well-defined functions in axon guidance and synapse formation in the nervous system, calcium-independent cell-cell adhesion in the immune system, and recognition in antigen and growth factor binding in signal transduction pathways (16–19). Recent work from the sponge and the fruit fly highlighted the importance of IgCAM proteins in multicellular development and epithelial integrity (20–22).

Building on these findings, results from knockdown experiments reported in this work underscore the importance of NCAM2 in *Nematostella* epithelial morphogenesis. Moreover, we found that embryos depleted for NCAM2 fail to gastrulate. Together, our results bring to light the important role of this neural cell adhesion molecule in both epithelial and germ layer organization during embryo development.

## Results

### Proteomic discovery of *Nematostella* embryo surface proteins

To identify cell surface proteins from *Nematostella* embryos, we treated unfertilized oocytes and early embryos with a non-permeable biotinylation reagent, Sulfo-NHS-LC Biotin, which labels cell surface proteins through its reaction with primary amines (23). Mitosis causes the early embryo epithelium to open up, so we arrested embryos in interface with a cell cycle inhibitor (8). After embryo lysis, biotinylated proteins were enriched by streptavidin binding and analyzed by mass spectrometry. We rank-ordered candidates by labeling efficiency and homology to known adhesion protein families, then selected the predicted transmembrane and membrane-associated proteins listed in Figure 1.

**Fig. 1.**
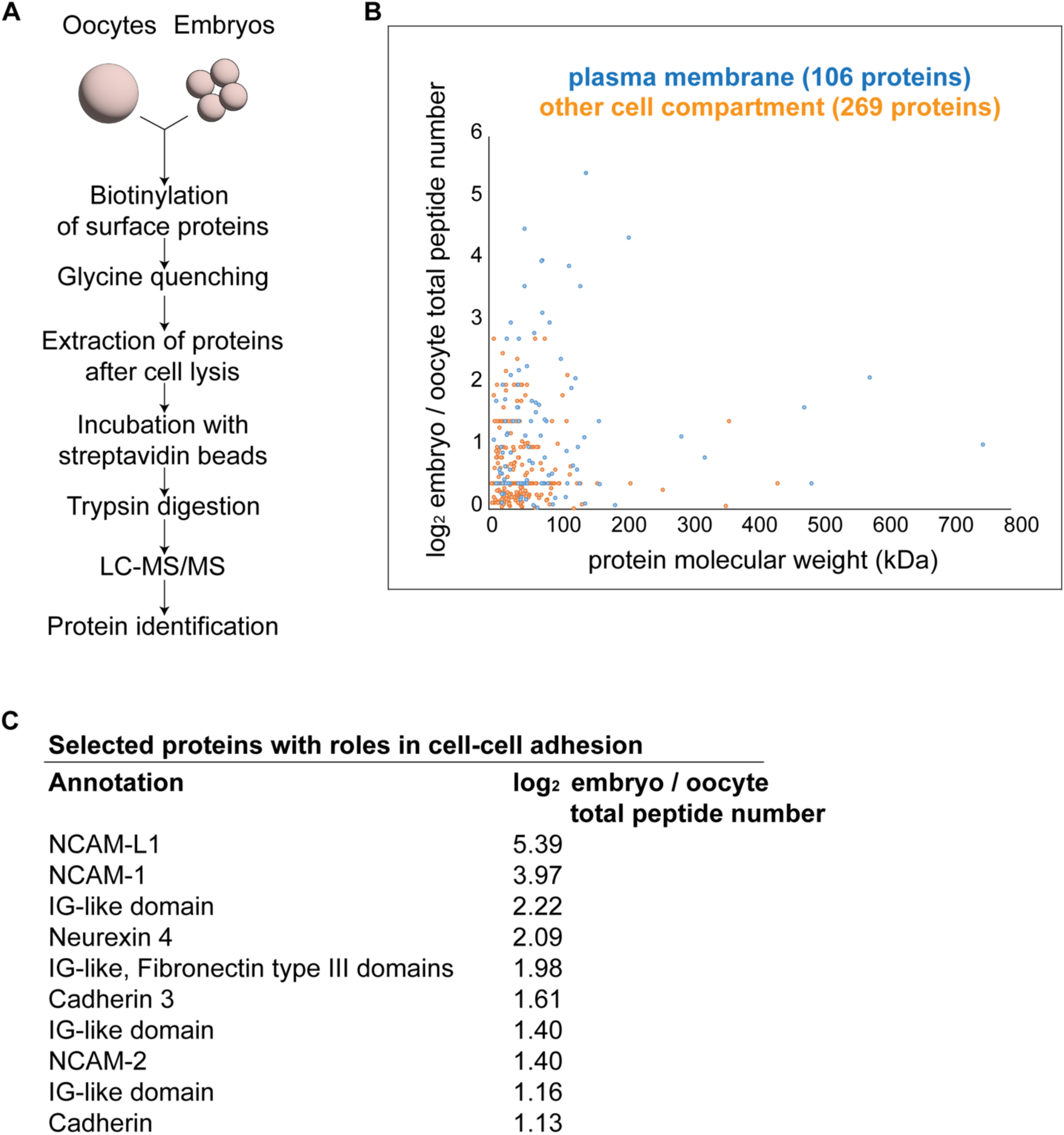
Experimental outline of cell surface protein labeling and discovery. (A) Workflow for protein labeling and peptide detection in *Nematostella* oocytes and embryos. (B) The numbers of total identified peptides from two independent experiments are plotted relative to the molecular weight of the corresponding proteins. Each dot depicts a predicted protein enriched in the plasma membrane (blue) or other intracellular compartment (orange). Only proteins with log_2_ ratios ≥ 0 are displayed. (C) The table shows the selected identified transmembrane proteins with known roles in cell-cell adhesion and their corresponding log_2_ ratios of identified peptides in embryos and oocytes.

Of the surface proteins detected, we found a good representation of the neural cell adhesion molecules that belong to the immunoglobulin superfamily (15). NCAM proteins mediate cell-cell adhesion and participate in signaling pathways that lead to cell differentiation (18, 24, 25). We also identified Cadherin3 which is expressed and localized in the ectoderm epithelium in early embryos, confirming the validity of our approach (13, 14). For our subsequent work, we focused on three NCAM proteins; NCAM1 and NCAM2 are predicted to contain five immunoglobulin-like (IgC2) and two fibronectin type III (Fn3) domains, while NCAML1 contains six IgC2 and five Fn3 domains (Fig. 2A).

**Fig. 2.**
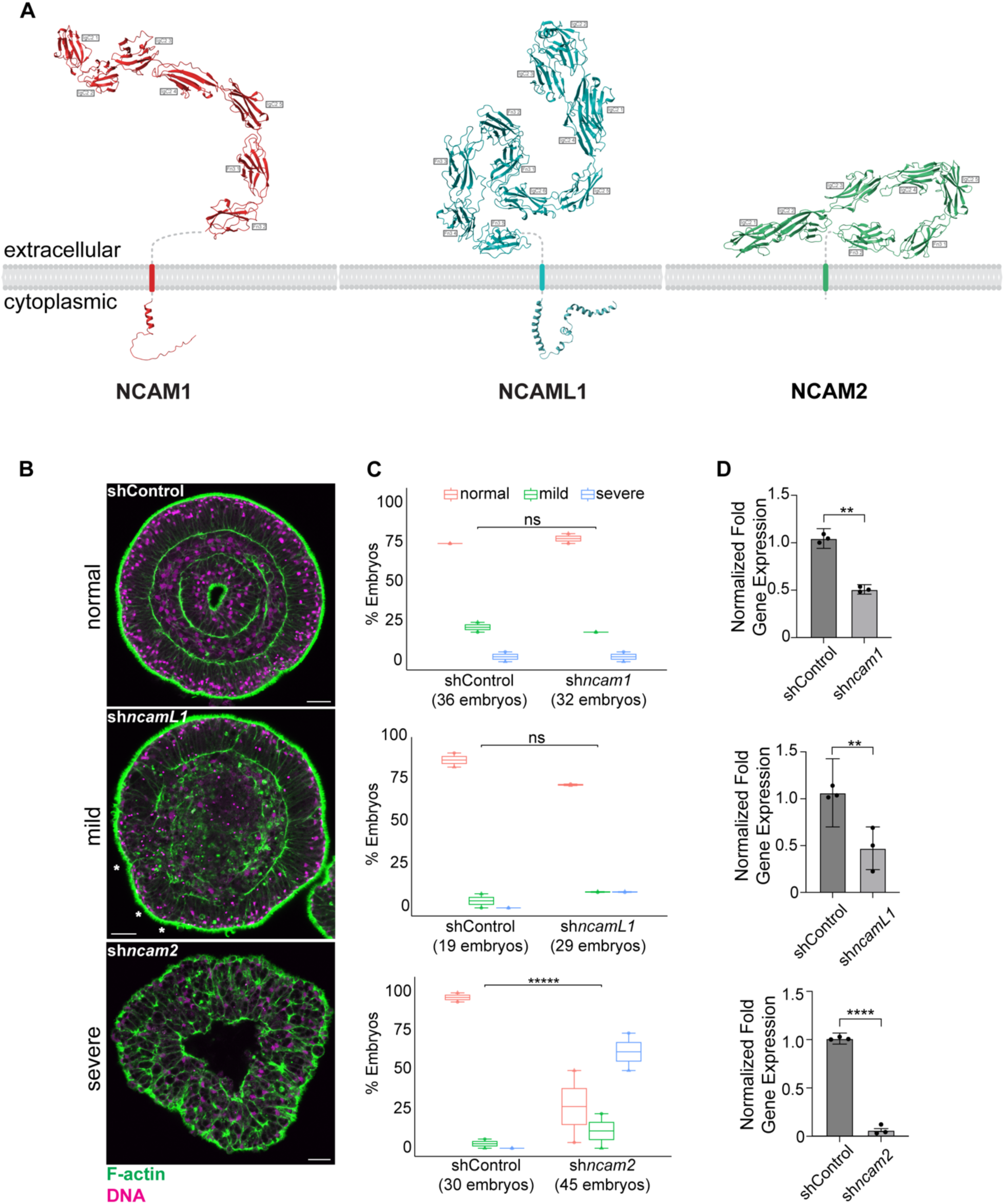
Depletion of NCAM2 impairs embryonic development. (A) Tertiary AlphaFold (v3) structure predictions of NCAM1, NCAML1 and NCAM2 (62). Extracellular and cytoplasmic domains were predicted separately and transmembrane domains are shown as solid bars. Immunoglobulin-like (IgC2) and fibronectin type III (Fn3) domains are indicated. (B) Representative midsagittal images of embryo phenotypes at gastrula stage showing clear outer and inner epithelial layers in the ‘normal’ category, regional disruptions in ‘mild’ embryos (indicated by the stars), or global disorganization with no distinct layers encountered in the ‘severe’ group. (C) Quantification of the three phenotypes, color-coded as indicated, in sh*ncam1*, sh*ncamL1* and sh*ncam2* knockdown embryos. Symbols correspond to embryo percentages from two independent experiments (Fisher’s exact test; ns: *p* > 0.05, \*\*\*\*\**p* < 0.0001). (D) Reduction of gene expression verified by qPCR of mRNA levels from embryos of three independent experiments (Wilcoxon rank sum test; \*\**p* = 0.0019, \*\*\*\**p* < 0.0001). Scale bar is 20μm.

### The neural cell adhesion molecule NCAM2 is required for embryonic development

To test if any of the NCAM family members from our proteomic screen were required for embryonic development, we individually knocked down NCAM1, NCAML1 and NCAM2 using RNA silencing and then assayed for epithelial organization and gastrulation efficiency. Gene expression examination by qPCR confirmed that the short RNA hairpins led to efficient mRNA knockdown. Of the three, knockdown of NCAM2 led to a high number of deformed embryos (Fig. 2B-D). By 26 hours post fertilization (hpf), embryos from the control group had completed gastrulation, and the inner and outer layers were clearly discernible. In sh*ncam2* knockdown embryos, the two layers were not distinguishable, suggesting either deformed epithelialization, failed gastrulation or both.

### NCAM2 is localized at apical cell junctions and is important for epithelial organization

To examine the subcellular localization of NCAM2, we first generated a specific polyclonal antibody against a peptide sequence present in a linker portion of its extracellular domain. We then tested the specificity of our NCAM2 antibody by examining its ability to detect and bind to the protein in control versus knockdown embryos. Gene expression examination by hybridization chain reaction (HCR) *in situs* confirmed that our short hairpin approach led to efficient *ncam2* mRNA knockdown (Fig. S1A-B). Both immunohistochemistry and immunoblot analysis of total cell extracts confirmed the antibody specificity and further verified the RNA knockdown efficiency. Immunoblot analyses from gastrula stage embryos showed significantly reduced NCAM2 levels but not complete elimination. Considering that NCAM2 was detected with our proteomic analysis in early embryos, we deduced that some of the protein may be maternally provided. Notably, based on its predicted molecular weight, the majority of the protein was detected in a likely dimer configuration (Fig. S1C).

Immunohistochemistry of embryos revealed that NCAM2 is localized at the apical cell junctions (Fig. 3A-B). The protein is localized at the apical junctions of the ectodermal epithelium, before and after gastrulation, with detectable signal at the apical junctions of the internalized layer (Fig. 3A, yellow arrows and Fig. S2A-B). We asked whether NCAM2 is co-localized with other known apical cell junction proteins such as β-catenin of the cadherin-catenin complex (13, 14, 26–28). NCAM2 and β-catenin co-localize at the apical junctions. However, of the two, only β-catenin is also enriched at the basal junctions (Fig. 3D and Fig. S2C green arrows). In sh*ncam2* embryos, NCAM2 junction enrichment was reduced. Importantly, the apical cell junctions as revealed by both NCAM2 and β-catenin were discontinuous and deformed in both late blastula and gastrula embryos (Fig. 3 and Fig. S2). While gastrula is the stage by which sh*ncam2* embryos show the most striking developmental defect, we examined embryos at the late blastula stage as well, when the ectodermal epithelium is fully mature but no morphogenetic events have been initiated yet.

**Fig. 3.**
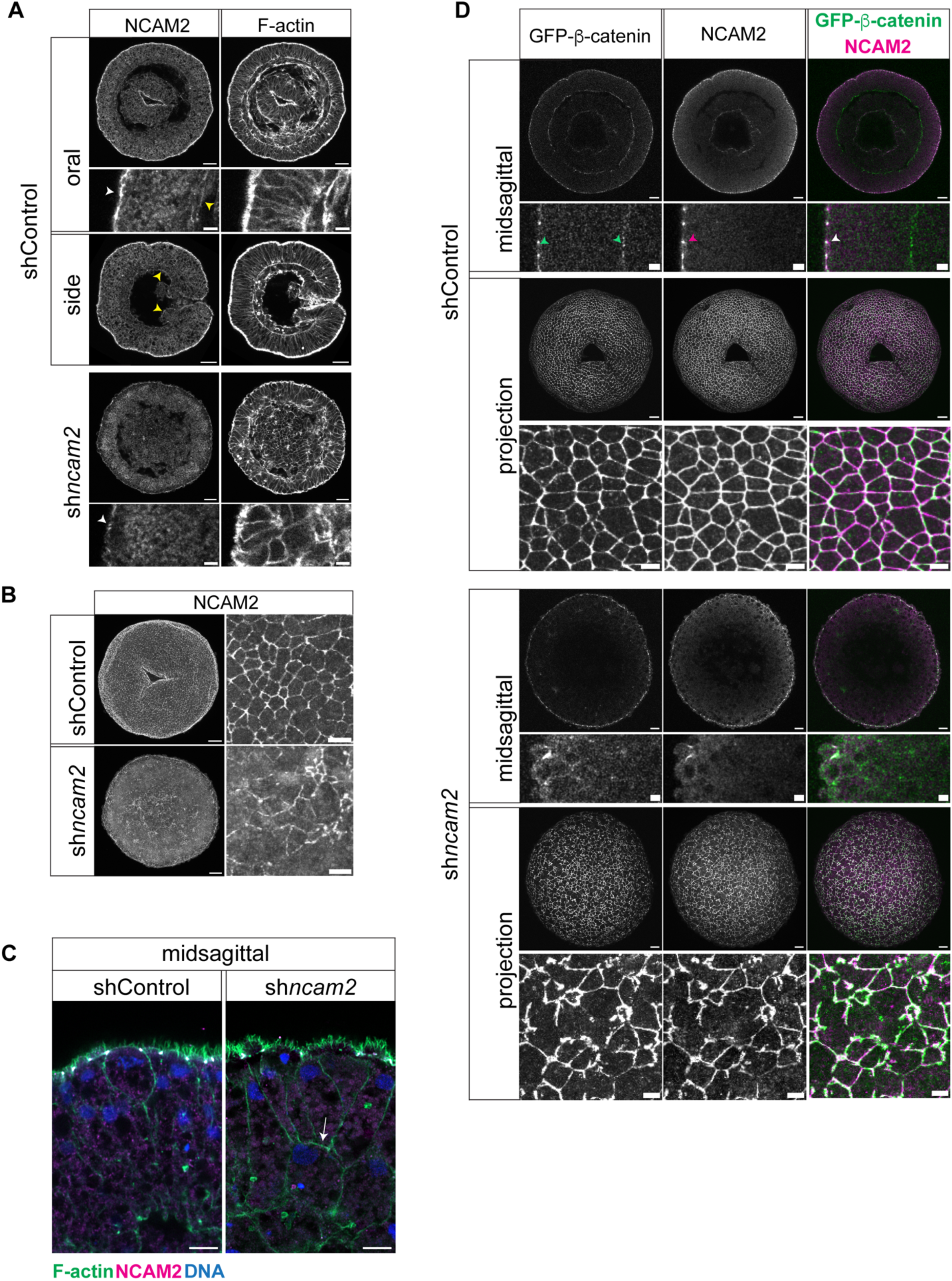
NCAM2 localizes at the apical cell junctions and is required for epithelial organization. (A) Oral and side views of midsagittal sections from gastrulating embryos showing NCAM2 localization at the apical junctions. White and yellow arrowheads point to the outer and inner epithelial apical junctions respectively. Blastopore is shown in the center and on the right of shControl embryos viewed from their oral end or side respectively. No distinct inner layer and no blastopore is visible in sh*ncam2* embryos. (B) Surface views of maximum projections showing reduced NCAM2 levels and apical junction disorganization in sh*ncam2* embryos. (C) Midsagittal view of a single cell layer in a shControl and a sh*ncam2* embryo in which an internalized cell is shown with no obvious NCAM2 detectable (internalized cell is indicated by the white arrow). (D) NCAM2 is co-localized with sfGFP-β-catenin at the apical cell junctions. Green arrowheads point to sfGFP-β-catenin localization at the apical and basal junctions; magenta arrowhead points to NCAM2 localization at the apical junctions and white arrowhead points to sfGFP-β-catenin and NCAM2 co-localization at the apical junctions. Projection views show reduced NCAM2 levels and apical junction disorganization in sh*ncam2* embryos as shown in (B). All close-up sections are cropped from the embryos shown above them with the apical domain shown on the left. Blastopore is evident in the center of shControl embryos in (B) and (D). Scale bar in close-up sections of (A) and (D) is 5μm and in all other images it is 20μm.

Our knockdown phenotype suggests that NCAM2 depletion leads to aberrant apical junctions. To test this hypothesis, we dissociated embryos at late blastula and gastrula stages and examined the ability of their cells to re-epithelialize after aggregation. *Nematostella* cells from dissociated embryos can readily organize by beginning to build an outer epithelium 6 hours post dissociation (hpd) (14, 26, 29). Compared to controls, dissociated cells from sh*ncam2* embryos failed to build a smooth apical domain even after 24hpd (Fig. 4). Eventually, all aggregates from sh*ncam2* embryonic cells disintegrated. From these experiments, we conclude that in the absence of NCAM2, cells fail to establish apical cell junctions and assemble an organized epithelium.

**Fig. 4.**
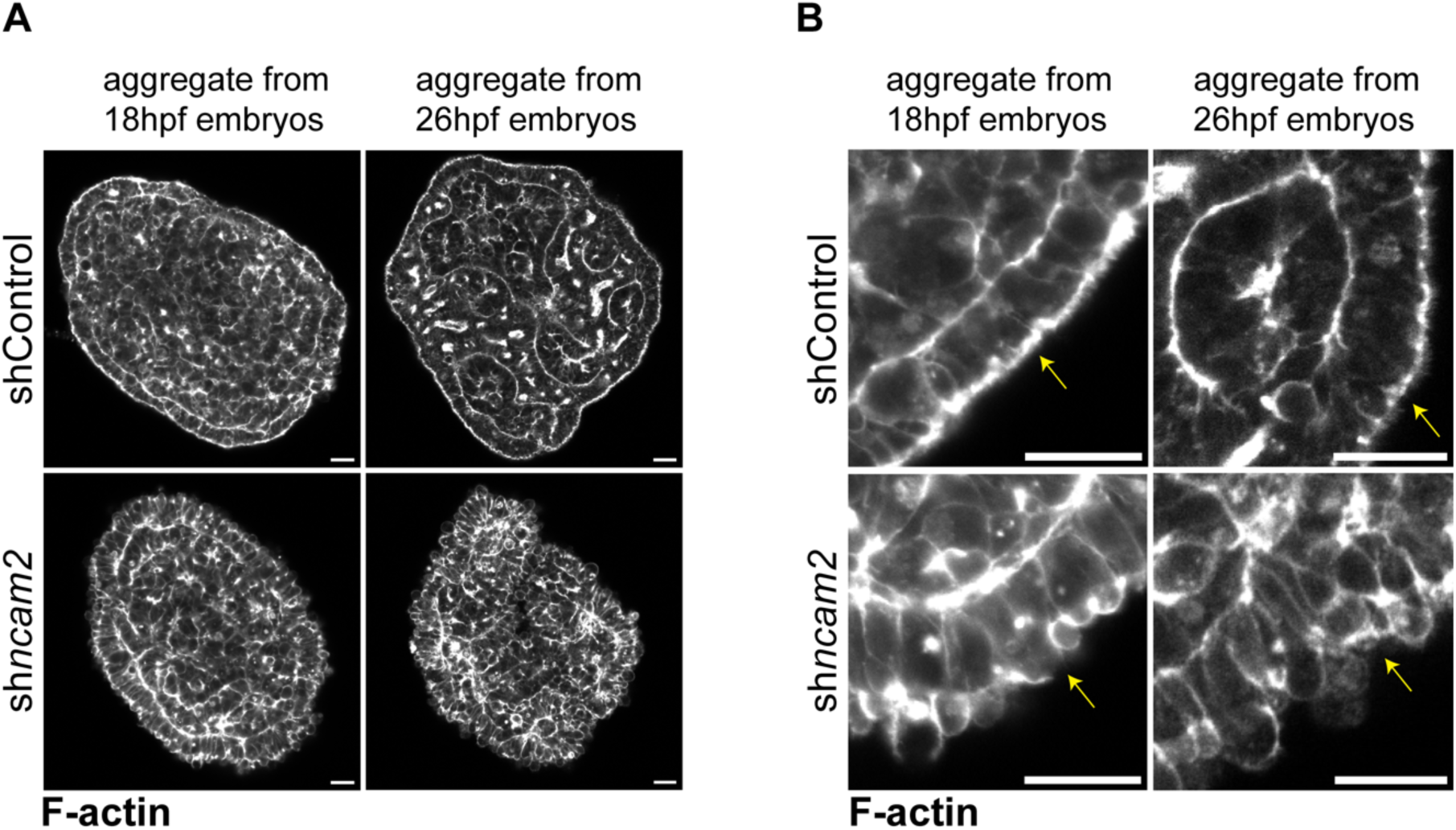
Cell aggregates from dissociated sh*ncam2* embryos fail to organize a smooth apical domain. (A) Aggregated cells were tested for epithelialization after dissociation of shControl and sh*ncam2* knockdown embryos at late blastula (18hpf) and gastrula stages (26hpf). Cells from sh*ncam2* embryos of both stages failed to form an outer layer. (B) Close-up sections show that cells from the sh*ncam2* aggregates fail to organize a smooth apical domain (pointed by the yellow arrows in all images). Scale bar is 20μm.

Interestingly, in midsagittal sections of the ectoderm epithelium, round cells were observed within the monolayer with no detectable NCAM2 on their surface (Fig. 3C). At this point, we hypothesized that as new cells arise in developing embryos they fail to integrate within the epithelium, likely due to insufficient NCAM2 levels. The gradual accumulation of non-epithelial cells failing to form apical junctions could compromise the integrity of the monolayer overall, with imposed strain on the junctions of neighboring epithelial cells. If this holds true, we would predict that sh*ncam2* embryos manifest disruptions in their basal cell junctions as well.

### Epithelial weakening in sh*ncam2* knockdown embryos

In late blastula embryos that were depleted of NCAM2, many cells appeared to extrude from the basal side of the epithelium (Fig. 5A). It is not clear whether these cells are actively ‘escaping’ the epithelium or are passively ‘removed’ from the epithelium because of loose cell adhesions. To examine the integrity of basal adhesions, we checked the localization of Cadherin3, as well as the phosphorylated myosin light chain (PMLC) at the basal domain of the ectodermal epithelium (Fig. 5B-C and Fig. S3). Both Cadherin3 and PMLC enrichment reveal the adhesion and high contractility at the apical and basal cell junctions (30). We scored quadrant sections of embryos and found significantly less Cadherin3 basal enrichment in sh*ncam2* embryos. This pattern was observed in both late blastula and gastrula stage embryos (Fig. 5B). Similarly, PMLC enrichment revealed a similar pattern (Fig. S3). Taken together, these results suggest that the basal epithelial domain is disrupted in sh*ncam2* embryos.

**Fig. 5.**
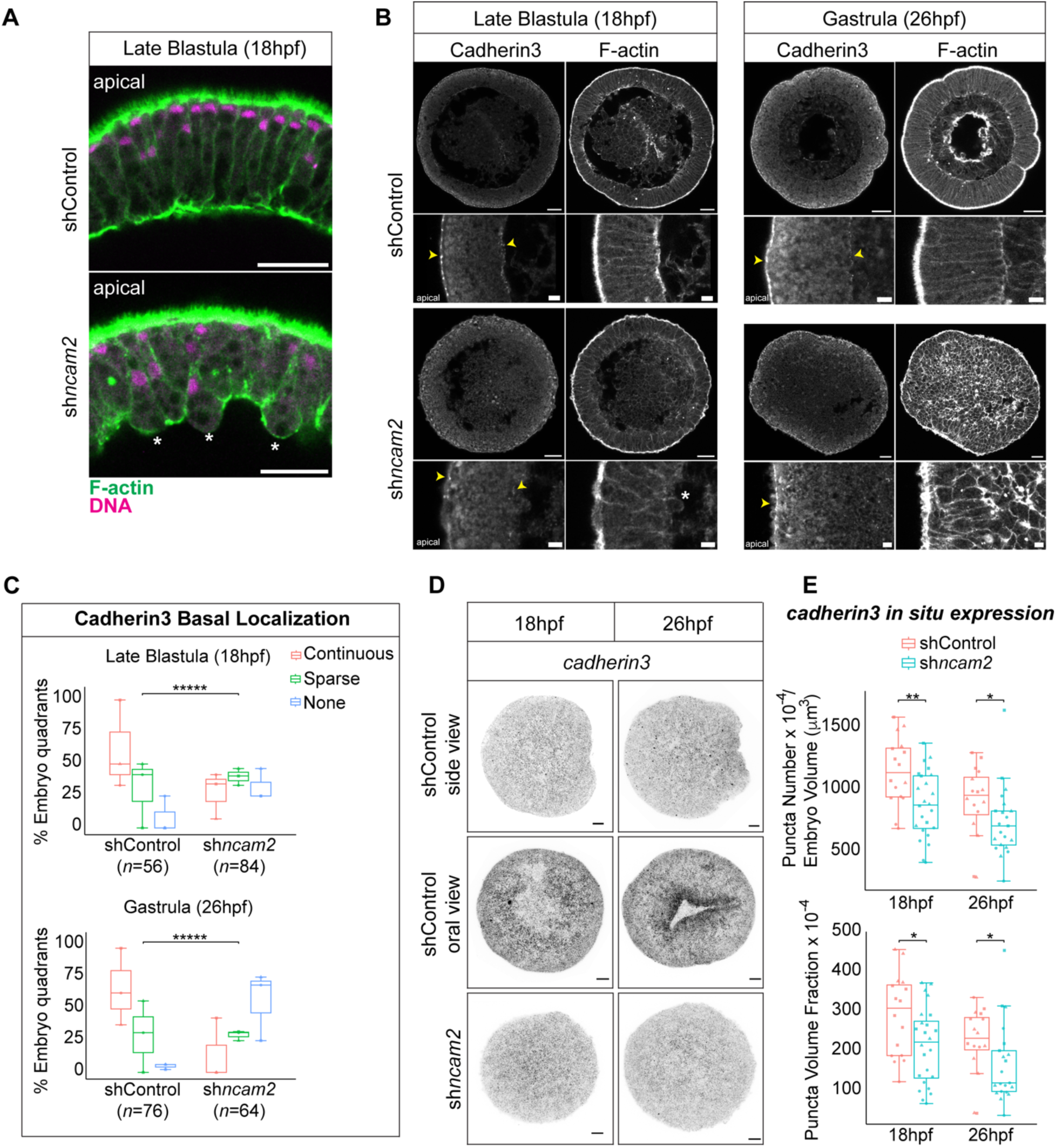
The basal epithelial domain is disorganized in sh*ncam2* knockdown embryos. (A) Midsagittal view of the epithelial monolayer at late blastula stage showing cell extrusion in sh*ncam2* knockdown embryos. Stars indicate basally extruding cells. (B) Midsagittal views showing Cadherin3 localization at the apical and basal junctions in late blastula and gastrula stage embryos. Yellow arrowheads point to apical and basal junctions. (C) Quantification of Cadherin3 basal localization in embryo quadrant sections, color-coded red for continuous, green for sparse and blue for none detectable. Symbols correspond to embryo quadrants from three independent experiments (Fisher’s exact test; \*\*\*\*\**p* < 0.0001). (D) *cadherin3* mRNA levels detected *in situ* in late blastula and gastrulating embryos. Due to lack of a definitive blastopore in sh*ncam2* embryos, their orientation is not determined. (E) Quantification of *cadherin3* mRNA levels by puncta counting per sampled embryo volume and puncta area fraction. Symbols correspond to embryo percentages from three independent experiments (Wilcoxon rank sum test; \**p* < 0.05, \*\**p* = 0.006). Number of shControl embryos analyzed per time point: 18hpf (16 embryos) and 26hpf (16 embryos). Number of sh*ncam2* embryos analyzed per time point: 18hpf (24 embryos) and 26hpf (21 embryos). Blastopore is shown in the center of gastrulating shControl embryos in (B) and (D). Scale bar in close-up sections of (B) is 5μm and in all other images it is 20μm.

Cell extrusions concurrent with disruption of the basal domain of the ectodermal epithelium in sh*ncam2* embryos are suggestive of epithelial-mesenchymal transitions (EMT) that occur during gastrulation in many species (31). In *Nematostella*, these transitions are constrained and partially occur at the invaginating plate of gastrulating embryos (32, 33). However, sh*ncam2* embryos fail to gastrulate as evidenced by the lack of a bi-layered organization. We searched for molecular signatures of potential EMT occurring in sh*ncam2* embryos by checking the expression patterns of genes widely shown to change when cells lose their epithelial identity. Down-regulation of cadherin is one of the hallmarks of EMT (34–37). We examined *cadherin3* mRNA levels by *in situ* HCR and found a significant reduction in both late blastula and gastrula embryos (Fig. 5D-E). While reduced NCAM2 and Cadherin3 levels could explain the disrupted apical and basal junctions of the epithelium, they do not adequately explain the all-encompassing disruption of development seen in sh*ncam2* embryos. The zinc-finger transcription factor Snail promotes EMT by repressing expression of cadherins and other proteins required for epithelial integrity (35, 37, 38). In *Nematostella*, *snailA* expression is confined to the invaginating plate of the embryo during gastrulation (7, 32, 39, 40). However, in sh*ncam2* embryos *snailA* was ectopically expressed throughout the ectodermal epithelium (Fig. 6A-B). This observation is consistent with widespread disruption of the epithelial cell state and initiative of an EMT program when NCAM2 is depleted.

**Fig. 6.**
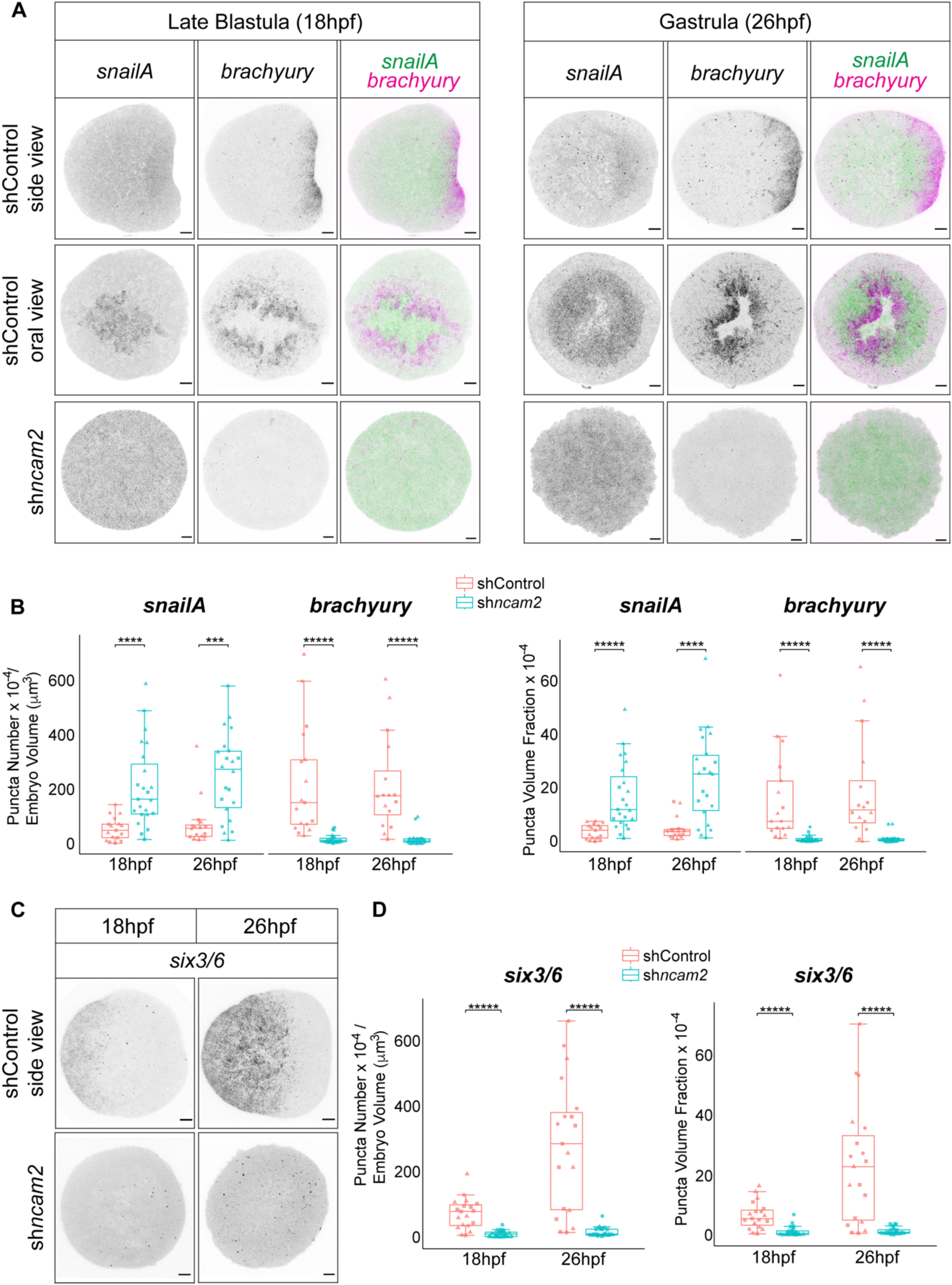
Germ layer identity is perturbed in embryos with reduced NCAM2 levels. (A) *In situ* detection of both *snailA* and *brachyury* mRNA levels in late blastula and gastrulating embryos. Ectopic expression of *snailA* and down-regulation of *brachyury* in sh*ncam2* embryos indicate aberrant germ layer identity. Blastopore is shown in the center of gastrulating shControl embryos viewed from their oral domain. (B) Quantification of *snailA* and *brachyury* mRNA levels by puncta counting per sampled embryo volume and puncta area fraction. Both *snailA* and *brachyury* were examined in the same embryos. Symbols correspond to embryo percentages from three independent experiments (Wilcoxon rank sum test; \*\*\**p* < 0.001, \*\*\*\*\**p* and \*\*\*\**p* < 0.0001). Number of shControl embryos analyzed per time point: 18hpf (17 embryos) and 26hpf (16 embryos). Number of sh*ncam2* embryos analyzed per time point: 18hpf (23 embryos) and 26hpf (22 embryos). (C) Aboral detection of *six3/6* mRNA levels in late blastula and gastrulating shControl embryos. Down-regulation of *six3/6* in sh*ncam2* embryos suggests defects in whole embryo patterning. (D) Quantification of *six3/6* mRNA levels by puncta counting per sampled embryo volume and puncta area fraction. Symbols correspond to embryo percentages from three independent experiments (Wilcoxon rank sum test; \*\*\*\*\**p* < 0.0001). Number of shControl embryos analyzed per time point: 18hpf (19 embryos) and 26hpf (19 embryos). Number of sh*ncam2* embryos analyzed per time point: 18hpf (23 embryos) and 26hpf (21 embryos). Scale bar is 20μm.

### Germ layer identity is perturbed in embryos with reduced NCAM2 levels

Expression of specific transcription factors demarcate the distinct territories occupied by the germ layers as they form during gastrulation. Recent work has re-defined the germ layers of *Nematostella* embryos. The inner layer that arises with invagination is homologous to mesoderm, while the blastopore ectoderm is more like the endoderm of bilaterians (41, 42). Accordingly, *snailA* expression is restricted to the mesoderm germ layer. Ectopic *snailA* in sh*ncam2* embryos prompted us to examine expression of other germ-layer specific factors. The blastopore lip of control embryos is marked by expression of the pre-endoderm-specific factor *brachyury* (Fig. 6A-B) (32, 43). In sh*ncam2* embryos, *brachyury* expression was barely detectable. Notably, *brachyury* expression at the blastopore lip has been correlated with the process of invagination and bending of the tissue during gastrulation (44, 45). It is possible that in the lack of *brachyury* expression, sh*ncam2* embryos fail to initiate invagination and gastrulate. The germ layer identity was also perturbed in other regions of the embryo, away from the mutually exclusive *brachyury*-*snailA* zone near the blastopore. As such, the aboral ectoderm-specific expression of *six3/6* was also abolished in sh*ncam2* embryos (Fig. 6C-D) (46). Together, these results show that NCAM2 depletion causes defects in whole-embryo patterning.

## Discussion

Comparative genomics have provided a list of conserved proteins that are essential for epithelial organization across Metazoa. With functional studies possible in early branching animals, like in *Nematostella*, we are able to test the role of many conserved proteins and probe the ancient origins of epithelial development. However, epithelialization is a context-dependent process, and an unbiased identification of epithelial proteins is especially important. In this study, we used biotinylation labeling and mass spectrometry to selectively identify surface proteins from early *Nematostella* embryos. We identified surface proteins that were previously detected in whole embryo proteomic analysis, including the early expressing Cadherin3 orthologue (47). Interestingly, we detected peptides from NCAM-like proteins, suggesting their presence on the surface of embryonic epithelia, well before the appearance of specialized neurons . NCAM proteins contain immunoglobulin-like and fibronectin type III domains participating in both intercellular adhesion and signaling functions in various tissues. Moreover, recent work has highlighted the importance of these proteins in sponge development and fly epithelial homeostasis supporting their ancient roles in allorecognition and cell adhesion (20–22). We discovered that NCAM2 is required for *Nematostella* development through its role in two important processes: 1) epithelial organization and 2) gastrulation.

We found NCAM2 to co-localize with β-catenin at the apical cell adhesions. Even though it is tempting to speculate a potential role of NCAM2 in the cadherin-catenin complex, it is intriguing that their co-localization only occurs apically. In *Nematostella*, Cadherin3, α-catenin, and β-catenin are enriched at both apical and basal junctions (14, 26). If NCAM2 functions together with these proteins, its activity is restricted to the apical domain. Indeed, our work suggests that NCAM2 is important for the integrity of apical cell junctions. In sh*ncam2* knockdown embryos the apical junctions appear disrupted, and in epithelialization aggregates the apical domain fails to organize. Eventually, both embryos and aggregates disintegrate suggesting severe epithelialization defects. We also found disrupted basal junctions. In embryos, Cadherin3 basal localization was compromised and cells appeared to undergo basal extrusion. Together, these manifestations suggest that NCAM2 is required for epithelial organization in *Nematostella*. How is its epithelial role mediated? One possibility is that NCAM2 is actively involved in cell-cell adhesions. The immunoglobulin-like domains have been shown to engage with each other in cis- and trans-configurations (48). AlphaFold (v3) structural predictions using the *Nematostella* protein sequence suggest a potential NCAM2-NCAM2 dimer formation via an interaction between the immunoglobulin-like domain of one monomer and the fibronectin type III domain of another (Fig. S4). However, this is rarely encountered and the predicted score is of low confidence (Predicted TM-score (pTM) = 0.39). Future work can specifically examine the adhesive function of NCAM2 and dissect the role of its domains.

We were most surprised to find that sh*ncam2* embryos fail to gastrulate. One can argue that a deformed epithelium fails to undergo the proper morphogenetic changes required for tissue invagination during gastrulation. Indeed, cells at the blastopore region require intact and well polarized domains to constrict apically and expand their basal domains (32, 33, 49). Interfering with myosin contraction blocks gastrulation, underscoring the importance of epithelial architecture in the mechano-transduction pathways active during the process (45, 50–52). Does depletion of other known epithelial proteins disrupt *Nematostella* gastrulation? Knockdown of Cadherin3 blocks gastrulation movements and depletion of α-catenin leads to endo-mesoderm defects after gastrulation (13, 14). These variable gastrulation outcomes may be due to differences in the efficacy of knockdown approaches or in the severity of epithelial defects.

Importantly, depletion of NCAM2 led to germ layer patterning defects. The ectopic expression of *snailA* suggests an activation of an epithelial-to-mesenchymal program (35, 37, 38). This event could be sufficient to disrupt global patterning. Given that *Nematostella* embryos gastrulate primarily by tissue folding, ectopic EMT in the absence of NCAM2 disrupts epithelialization that precedes gastrulation and further development. Morphant embryos with blocked β-catenin expression share a similar phenotype with sh*ncam2* embryos in both their failed gastrulation and germ layer patterning defects (28). NCAM2 as a surface protein could directly or indirectly impact on signaling pathways that intersect with β-catenin. It is worth noting that the predicted NCAM2 protein structure does not include a cytoplasmic domain (Fig. 2A). Any effect of NCAM2 onto other signaling molecules likely occurs through extracellular interactions. Alternatively, NCAM2 does not directly participate in any signaling pathways. Instead, signaling pathways best function within well-organized epithelia. Testing of all these hypotheses is currently under way.

## Acknowledgments

We are grateful to Ross Tomaino (Taplin Mass Spectrometry Facility, Harvard Medical School) for mass spectrometry and data acquisition, Grigory Genikhovich (University of Vienna) for the sfGFP-β-catenin transgenic animals, Elizabeth Van Itallie (Harvard Medical School) for help with protein extraction, Lampros Panagis (Amherst Biology Imaging Center) for microscopy training, advice and helpful discussions, Aissam Ikmi (EMBL) for help with HCR, Amira Abdelghany for help with HCR quantification, Caroline Towse and Lauren Wagner for help with animal maintenance and Leah Davis for administrative support. We acknowledge the generous gift of Cadherin3 antibody from Ulrich Technau (University of Vienna) which was unfortunately damaged during shipping. We also thank the members of the Mitchison lab for their help and hospitality during K.R.’s sabbatical visit. This work was supported by Amherst College, a National Science Foundation grant MRI-2117798 and a National Institute of General Medical Sciences grant R15GM141979-01.

## Materials and Methods

### Animal and embryo handling

*Nematostella vectensis* animals were cultured in 12 parts per thousand (ppt) artificial sea water (SW) (Sea Salt: Instant Ocean) at 17°C, were fed weekly with freshly hatched *Artemia salina* brine shrimp larvae and were spawned every 3 weeks (53). Oocytes were de-jellied soon after spawning and were electroporated with shRNA as previously described (54). Treated oocytes were fertilized immediately after electroporation and embryos were kept at 21°C.

### Biotinylation and isolation of surface proteins from oocytes and embryos

Oocytes and cell-cycle arrested embryos (for drug treatment see below) were incubated at room temperature (RT) for 10min with 5mM EZ-Link Sulfo-NHS-LC Biotin (Thermo Scientific, A39257) in PBS pH 7.4 (23). Residual biotin was quenched by washing the samples once with 20mM glycine in PBS pH 7.4 for 3min. Excess buffer was removed and samples were snap frozen in liquid nitrogen and stored at -80°C. Frozen samples were treated on ice for 10min with lysis buffer (25mM HEPES (pH 7.2), 250mM sucrose, 1% Nonidet P-40 Alternative (Sigma 492016), 10mM EDTA (pH 7.2), EDTA-free mini protease inhibitor (Pierce 11836170001), 10μM Combretastatin 4A and 10μM Cytochalasin D) as described previously (55). The cytoplasmic and lipid layers were separated from the pellet after a 4min centrifugation at 4,000 relative centrifugal force (RCF) at 4°C. The oocyte / embryo lysate was then incubated with streptavidin magnetic beads (Pierce 88816) with overnight rocking at 4°C. The bead / protein sample was washed twice with RIPA buffer (Pierce 89900) and five times with 50mM ammonium bicarbonate. The washed bead / protein sample was resuspended in 50mM ammonium bicarbonate before preparation for mass spectrometry.

### Mass spectrometry and data acquisition

The bead / protein samples were spiked with 10ng/μl modified sequencing-grade trypsin (Promega) and were placed in a 37°C room overnight. The samples were then placed on a magnetic plate and the liquid removed. The extracts were dried in a speed-vac (∼1 hr). Samples were then re-suspended in 50μl of HPLC solvent A (2.5% acetonitrile, 0.1% formic acid) and desalted by STAGE tip (56). On the day of analysis the samples were reconstituted in 10µl of HPLC solvent A. A nano-scale reverse-phase HPLC capillary column was created by packing 2.6µm C18 spherical silica beads into a fused silica capillary (100µm inner diameter x ∼30cm length) with a flame-drawn tip (57). After equilibrating the column each sample was loaded via a Famos auto sampler (LC Packings) onto the column. A gradient was formed and peptides were eluted with increasing concentrations of solvent B (97.5% acetonitrile, 0.1% formic acid). As peptides eluted, they were subjected to electrospray ionization and then entered into a Velos Orbitrap Elite ion-trap mass spectrometer (Thermo Fisher). Peptides were detected, isolated, and fragmented to produce a tandem mass spectrum of specific fragment ions for each peptide. Peptide sequences (and hence protein identity) were determined by matching protein databases with the acquired fragmentation pattern by the Sequest software program (Thermo Fisher) (58). All databases include a reversed version of all the sequences and the data were filtered to between a one and two percent peptide false discovery rate. For the total peptide display in Figure 1, the average of total peptide numbers quantified for a given protein from two independent experiments was normalized to the average number of carboxylase peptides and then the log_2_ ratio of embryo / oocyte peptides was plotted relative to the predicted protein molecular weight.

### Drug treatments

Embryos were arrested in the fully compacted state at 3hpf with addition of 10µM Flavopiridol hydrochloride (Tocris Bioscience 3094) in SW supplemented with 1% DMSO (8). After 1h drug treatment at RT, embryos were washed once with PBS pH 7.4 before biotinylation described above.

### Generation of NCAM2 and Cadherin3 antibodies

To detect *Nematostella vectensis* NCAM2 and Cadherin3, polyclonal antibodies were raised in rabbits against chemically synthesized peptides. Both antibodies were validated by enzyme-linked immunosorbent assay (Eurogentec).

### Immunoblot analysis

Protein samples from 100-250 embryos were denatured at 95°C for 5min after adding 2xLaemmli sample buffer (4% SDS, 0.04% Bromophenol Blue, 120mM Tris-HCl pH 6.8, 0.2% glycerol, 10% β-Mercaptoethanol). Total protein extracts were loaded into a 4-20% pre-cast polyacrylamide gel (Bio-Rad, 4568094), then blotted and probed with rabbit NCAM2 primary antibody (1:100) and anti-rabbit Horseradish Peroxidase (HRP)-conjugated secondary antibody (1:10,000, Jackson ImmunoResearch, 111-035-003). Protein levels were normalized by striping blots and probing extracts with mouse α-tubulin primary antibody (1:1000, Sigma, T6074) and anti-mouse HRP-conjugated secondary antibody (1:10,000, Invitrogen, A10668). Chemiluminescence was detected with Clarity Western ECL substrate (Bio-Rad, 1705060), captured with Syngene GeneGnome XRQ and quantified using Fiji (ImageJ) software (59).

### Gene knockdown by shRNAs

shRNAs were designed and prepared as previously described (26, 54). Sequences of genes of interest were checked by BLASTn against the *Nematostella vectensis* genome and after confirming specificity were used for primer design and ordering from Integrated DNA Technologies. shRNAs against *gfp* and *mcherry* were used as negative controls (shControl). Oocytes were electroporated with shRNAs (500ng/μl in 15% Ficoll PM400/SW) (26, 54).

### Quantification of gene expression

Knockdown of gene expression was confirmed by qPCR from three biological replicates as described before (26). Briefly, total cDNA was prepared from purified RNA using the iScript cDNA Synthesis Kit (Bio-Rad) and quantification of expression was done by qPCR that was carried out in three technical replicates for each sample, using the SsoAdvanced Universal SYBR Green Supermix (Bio-Rad) on a CFX Duet Real-Time PCR System (Bio-Rad). ATP synthase was chosen as the endogenous reference gene for normalization.

Gene expression was examined *in situ* by the Hybridization Chain Reaction (HCR) as described previously with some modification (60, 61). Probe sets were ordered from Molecular Instruments by using sequences with the following accession numbers; *cadherin3* (MK253652), *snailA* (AY651960)*, brachyury* (AF540387) and *six3/6* (KC137590) (14, 39, 46). Amplifiers, hybridization, wash and amplification buffers were ordered from Molecular Instruments, Inc (molecularinstruments.com). Embryos were fixed at 18hpf or 26hpf in ice-cold 4%PFA and 0.2% glutaraldehyde in SW on ice for 90sec and then in 4%PFA/SW at RT for 1h. After 5 washes in PTw (1xPBS, 0.1% Tween20), embryos were dehydrated through a PTw/Methanol gradient and stored in 100% Methanol at -20°C. After rehydration through a Methanol/PTw gradient, embryos were treated with 20μg/ml Proteinase K (Sigma P2308) in PTw for 5min at RT, washed 2 times with 4mg/ml glycine in PTw and 2 times with PTw, and fixed with 4% PFA/PTw for 30min at RT. After 3 washes with PTw, embryos were incubated in Probe Hybridization Buffer for 30min at 37°C. Embryos were then incubated overnight at 37°C with 1pmol probe set in 100ul Probe Hybridization Buffer. Embryos were then washed 2 times with the Probe Wash Buffer at 37°C for 30min, and after 2 washes with 5xSSCTw (Saline-Sodium Citrate buffer, 0.1%Tween20) at RT they were incubated with the Amplification Buffer for 1h at RT. Hairpins h1 and h2 (3pmol each) were heated to 95°C for 90sec, cooled to RT for 30min and were added to 100ul Amplification Buffer together with embryos for an overnight incubation at RT. Embryos were washed with 5xSSCTw, stained with 1μg/ml DAPI (Sigma D9542) and mounted for imaging.

### Embryo dissociation and cell aggregation

Embryos were dissociated at either 18hpf or 26hpf and their cells aggregated as previously described (26, 29). Briefly, embryos were collected in 100μl SW and were dissociated after addition of 200μl of Ca^2+^/Mg^2+^-free artificial sea water (27g/L NaCl, 1g/L Na_2_SO_4_, 0.8g/L KCI, 0.18g/L NaHCO_3_ in Milli-Q water). Cells were strained and aggregated via centrifugation for 30min at 2.700 x *g* (Sorvall Legend Microcentrifuge 21). Experiments were done at least three times for both time points.

### Fixation and immunohistochemistry

For DNA and F-actin staining, *Nematostella* embryos were fixed in 4%PFA and 0.2% glutaraldehyde in PBS for 90sec at RT and then overnight in 4%PFA/PBS at 4°C as previously described (26).

For all antibody staining procedures, embryos were fixed in 4% PFA/PBS for 1h at 4°C, followed by a 7min-incubation on ice with ice-cold acetone. After six washes in PBS/0.2% TritonX-100, embryos were incubated in blocking solution (PBS/0.2% TritonX-100, 5% normal goat serum (Jackson Immuno Research Labs), 1% bovine serum albumin (BSA), 1% DMSO) for 2h at RT before overnight incubation in the same buffer with the selected primary antibodies at 4°C. Primary antibodies were used in the following concentrations: rabbit NCAM2 (1:100), rabbit Cadherin3 (1:100), rabbit anti-GFP (1:500, abcam ab290) and mouse phospho-myosin light chain 2 (Ser19) antibody (1:200, Cell Signaling 3675). After six washes in PBS/0.2% TritonX-100, embryos were incubated with blocking solution for 2h at RT, followed by overnight incubation in the same solution with the chosen secondary antibodies at 4°C. The following secondary antibodies were all used in 1:1000 concentration: goat anti mouse Alexa Fluor 488 antibody (Invitrogen, A11001), goat anti-mouse Alexa Fluor 568 antibody (Invitrogen, A11004), goat anti-rabbit Alexa Fluor 488 (Invitrogen, A11008) and goat anti-rabbit Alexa Fluor 555 (Invitrogen, A21428). DNA was stained with either DAPI (1μg/ml) or Draq5 (1:2000, Cell Signaling 4084) and F-actin was stained with Alexa Fluor 488, 555 or 647 phalloidin (1:200, Invitrogen A12379, A34055 or A22287, respectively) that was added in the overnight secondary antibody incubation at 4°C. After two washes in PBS/0.2% TritonX-100, embryos were mounted in a 2:1 mixture of Benzyl Benzoate:Benzyl Alcohol.

For visualization of both sfGFP-β-catenin and NCAM2, oocytes from transgenic sfGFP-β-catenin animals (from Grigory Genikhovich, described in (27)) were electroporated with either shControl or sh*ncam2* and at 18hpf or 26hpf embryos were fixed in 4%PFA/PBS at 4°C for 1h. Then embryos were processed as described above, except that only the rabbit NCAM2 antibody was included in the primary incubation and embryos were mounted in Vectashield (Vector Laboratories) following a gradient of glycerol exchanges.

Aggregates were fixed in 4% PFA/PBS by overnight incubation at 4°C. Staining of DNA and F-actin, and mounting was performed as described above.

### Imaging

Fixed embryos and aggregates were imaged with a 40x oil objective (1.30NA) using an inverted confocal Nikon Ti Eclipse microscope, and with a 40x oil objective (1.40NA) on an inverted Zeiss LSM980 confocal microscope. Airyscan images on Figure 3C were obtained with an oil 63x High Strehl ratio objective (1.40NA) using the Zeiss LSM 980 microscope.

### Image analysis

Images were processed using Fiji (ImageJ) (59). Depending on the experiment, images were analyzed as follows:

For quantification of Cadherin3 or Phosphorylated Myosin Light Chain basal enrichment, 10μm-long Z sections encompassing the midsagittal part of embryos were maximally projected and divided into four XY area quadrants per embryo. Then quadrants were placed in one of three categories based on whether protein localization appeared continuous, sparse or none detected along the basal side of the epithelium.

Quantification of gene expression from HCR images was performed by calculating either the number or volume fraction of *in situ* hybridization puncta in the sampled volume of each embryo. To calculate the sampled embryo volume in μm^3^, the total area (μm^2^) of systematically randomly sampled sections spaced 1μm apart was obtained after segmentation and multiplied by the optical section thickness (0.6μm-0.8μm). Segmentation of hybridization puncta from the same optical sections was used to quantify the number and area of puncta. The quantification was expressed in terms of the total number of puncta in the sampled embryo volume (puncta number / embryo volume μm^3^) or as puncta volume fraction (sum of puncta area / sum of embryo section area). Debris or objects with area bigger than 2.5μm^2^ were computationally removed from the final analysis. HCR-treated embryos shown in figures are maximally projected with the exception of Figure S1A where one midsagittal section of an embryo is shown.

### AlphaFold predictions

The extracellular, transmembrane and cytoplasmic domains were identified via Phobius (*Phobius accessions: NON_CYTOPLASMIC_DOMAIN and SIGNAL_PEPTIDE*) and TMHMM (*TMHMM accession: TMhelix)*, and only the sequence of the extracellular region was retained unless stated otherwise. Domain composition of proteins was analyzed using InterProScan, with domain annotations referenced from the SMART member database. Two types of domains were identified; the immunoglobulin C2-set domain (*SMART accession: SM00408*) and the fibronectin type III domain (*SMART accession: SM00060*).

The NCAM1, NCAML1 and NCAM2 models were predicted with AlphaFold (v3), ranked by predicted template modeling (pTM) score and the highest-scoring model was selected for analysis (62). Raw output files, including the macromolecular crystallographic information (.cif) structure file and associated predicted aligned error (PAE) and predicted local distance difference test (pLDDT) scores (from *full_data_0* file) were downloaded and imported into UCSF ChimeraX (version 1.9) for structural analysis and visualization (63). Signal peptide regions were manually trimmed, referencing Phobius (*Phobius accession: SIGNAL_PEPTIDE*).

For the protein domain topology, the front-facing β-sheet was defined as the one containing the N-terminus, and the back-facing β-sheet as the one containing the C-terminus, and both were colored differently. Topology maps were drawn using arrows to represent β-strands and cylinders to represent α-helices, with the relative lengths and positions of secondary structure elements approximated from the predicted three-dimensional structures.

The interaction between NCAM2 – NCAM2 was modelled 40 times (25 using the extracellular region, 15 using the full sequence) using AlphaFold-Multimer (v3) with each job run using a unique random seed (62). Predicted models were ranked by predicted template modeling (pTM) score and interface-predicted template modeling-(ipTM) score. The highest-scoring model for each group was selected for further analysis. Additional high-confidence, lower-ranked models were examined to assess consistency of the predicted interaction patterns and rule out potential artifacts. For each highest-scoring model, raw output files were downloaded from AlphaFold-Multimer (v3) server. The highest-confidence internal model (*model_0*) macromolecular crystallographic information (.cif) structure file and its PAE and pLDDT scores (from *full_data_0* file) were loaded into ChimeraX for structural analysis and visualization (63). Molecular surfaces were rendered using the ChimeraX *surface* command. Predicted protein–protein interactions were visualized as contacts between the interacting residues using the *alphaFold contacts* command, which identifies residue pairs based on interatomic distance and predicted aligned error (PAE) thresholds.

### Statistical analysis

Data analysis and visualization was performed using Microsoft Excel, Prism 10 as well as R (4.5.1) *ggplot2* (3.5.2), *dplyr* (1.1.4), *purrr* (1.0.4), *ggpubr* (0.6.0), and *ggbeeswarm* (0.7.2) packages. Statistical analysis was performed via Wilcoxon rank sum exact test and Fisher’s exact test for count data. Significance level, alpha, was set to 0.05.

## Data Availability

The mass spectrometry proteomics raw data will be available via the PRoteomics IDEntification (PRIDE) partner repository upon final publication.

## Supplemental Figures

**Fig. S1.**
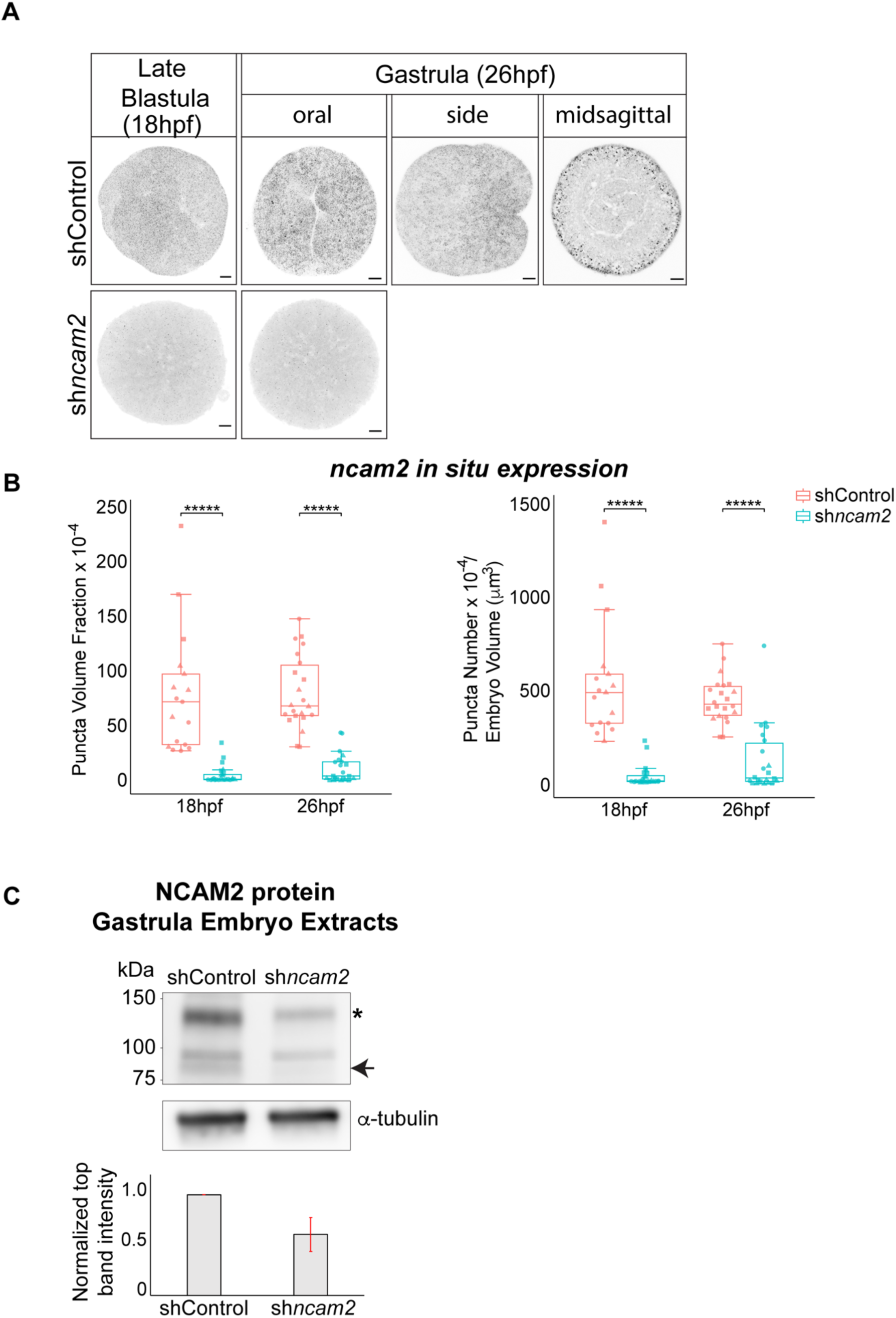
Validation of NCAM2 depletion in sh*ncam2* knockdown embryos. (A) *ncam2* mRNA levels detected *in situ* in late blastula and gastrulating embryos. Midsagittal view shows predominant *ncam2* expression in the outer epithelium. Blastopore is shown in the center of a gastrulating shControl embryo viewed from its oral domain. (B) Quantification of *ncam2* mRNA levels by puncta counting per sampled embryo volume and puncta area fraction. Symbols correspond to embryo percentages from three independent experiments (Wilcoxon rank sum test; \*\*\*\*\**p* < 0.0001). Number of shControl embryos analyzed per time point: 18hpf (17 embryos) and 26hpf (22 embryos). Number of sh*ncam2* embryos analyzed per time point: 18hpf (24 embryos) and 26hpf (22 embryos). (C) Immunoblot of NCAM2 from total protein extracts of gastrulating embryos. Star indicates potential NCAM2 dimer levels which appear reduced in the sh*ncam2* embryos. Arrow points to potential NCAM2 monomer that is not detectable in the sh*ncam2* embryo extract (predicted monomer molecular weight: 79kDa, predicted dimer molecular weight: 158kDa). Top band intensity levels were normalized to α-tubulin levels from the same embryo extracts. Top band levels from sh*ncam2* extracts were compared to top band levels from shControl extracts that were set to 1 for each immunoblot and the averaged value from three independent experiments was plotted. Error bar shows the standard deviation across experiments. Scale bar in images is 20μm.

**Fig. S2.**
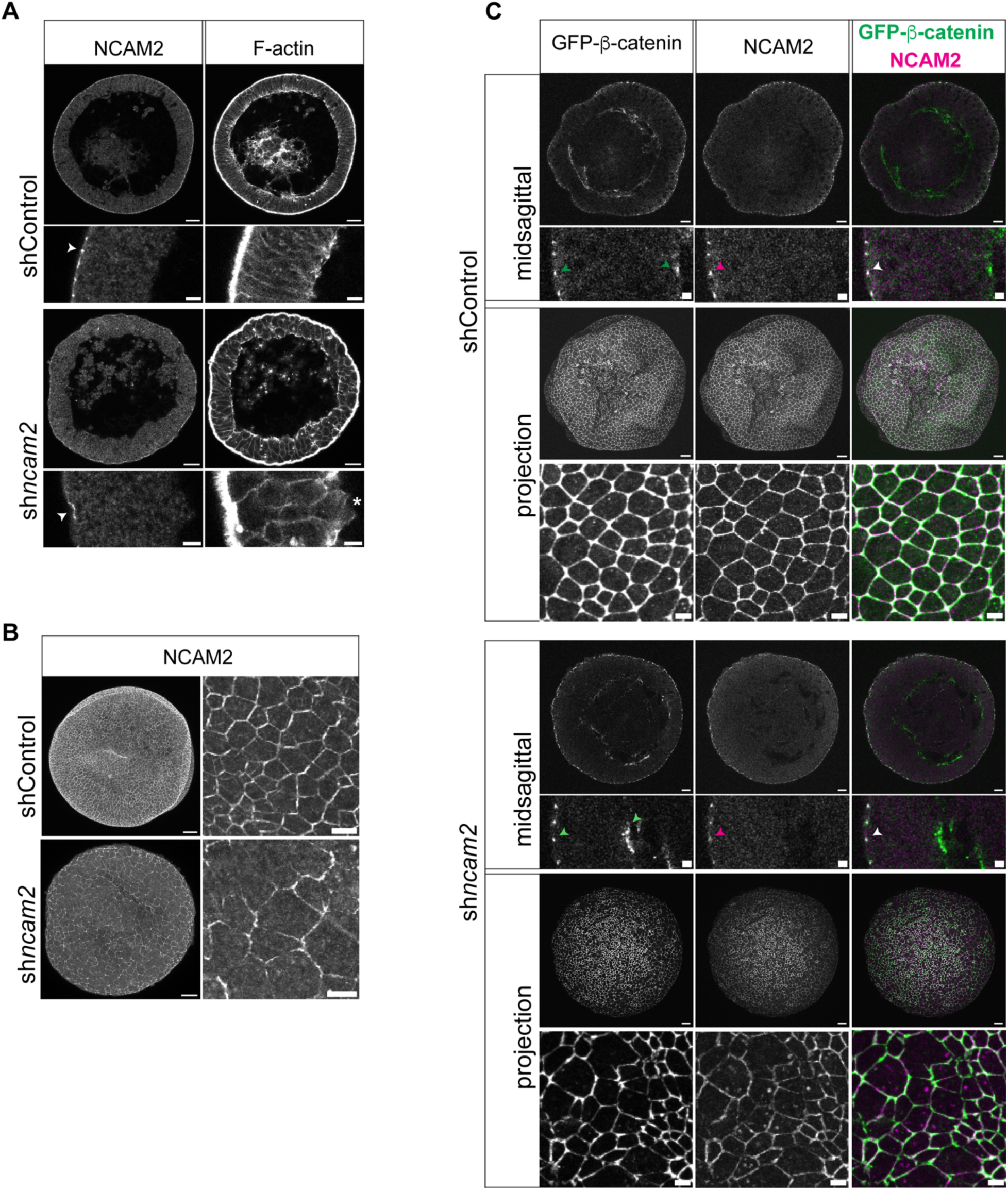
NCAM2 localizes at the apical cell junctions and is required for epithelial organization. (A) Side views of midsagittal sections from late blastula embryos showing NCAM2 localization at the apical junctions. White arrowheads point to apical junctions. Star indicates extruding cell. (B) Surface views of maximum projections showing reduced NCAM2 levels and apical junction disorganization in sh*ncam2* embryos. (C) NCAM2 co-localizes with sfGFP-β-catenin at the apical cell junctions. Green arrowheads point to sfGFP-β-catenin localization at the apical and basal junctions; magenta arrowhead points to NCAM2 localization at the apical junctions and white arrowhead points to sfGFP-β-catenin and NCAM2 co-localization at the apical junctions. Projection views show reduced NCAM2 levels and apical junction disorganization in sh*ncam2* embryos as shown in (B). All close-up sections are cropped from the embryos shown above them with the apical domain shown on the left. Scale bar in close-up sections of (A) and (C) is 5μm and in all other images it is 20μm.

**Fig. S3.**
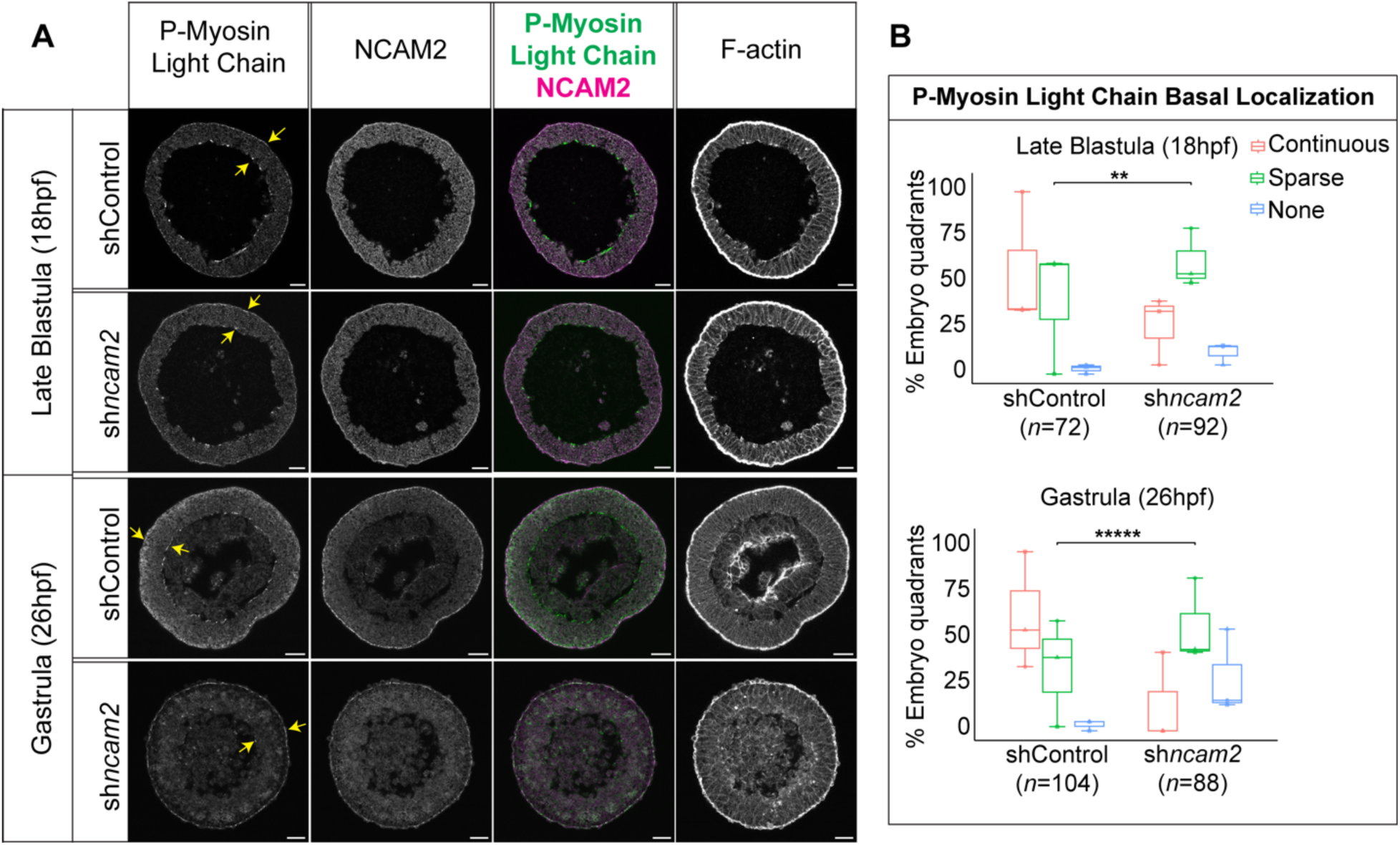
The basal epithelial domain is disorganized in sh*ncam2* knockdown embryos. (A) Midsagittal views showing phosphorylated myosin light chain (PMLC) enrichment at the apical and basal junctions in late blastula and gastrula stage embryos. Yellow arrows point to apical and basal junctions. Blastopore is shown in the center of a gastrulating shControl embryo. (B) Quantification of PMLC basal localization in embryo quadrant sections, color-coded red for continuous, green for sparse and blue for none detectable. Symbols correspond to embryo quadrants from three independent experiments (Fisher’s exact test; \*\**p* = 0.001, \*\*\*\*\**p* < 0.0001). Scale bar is 20μm.

**Fig. S4.**
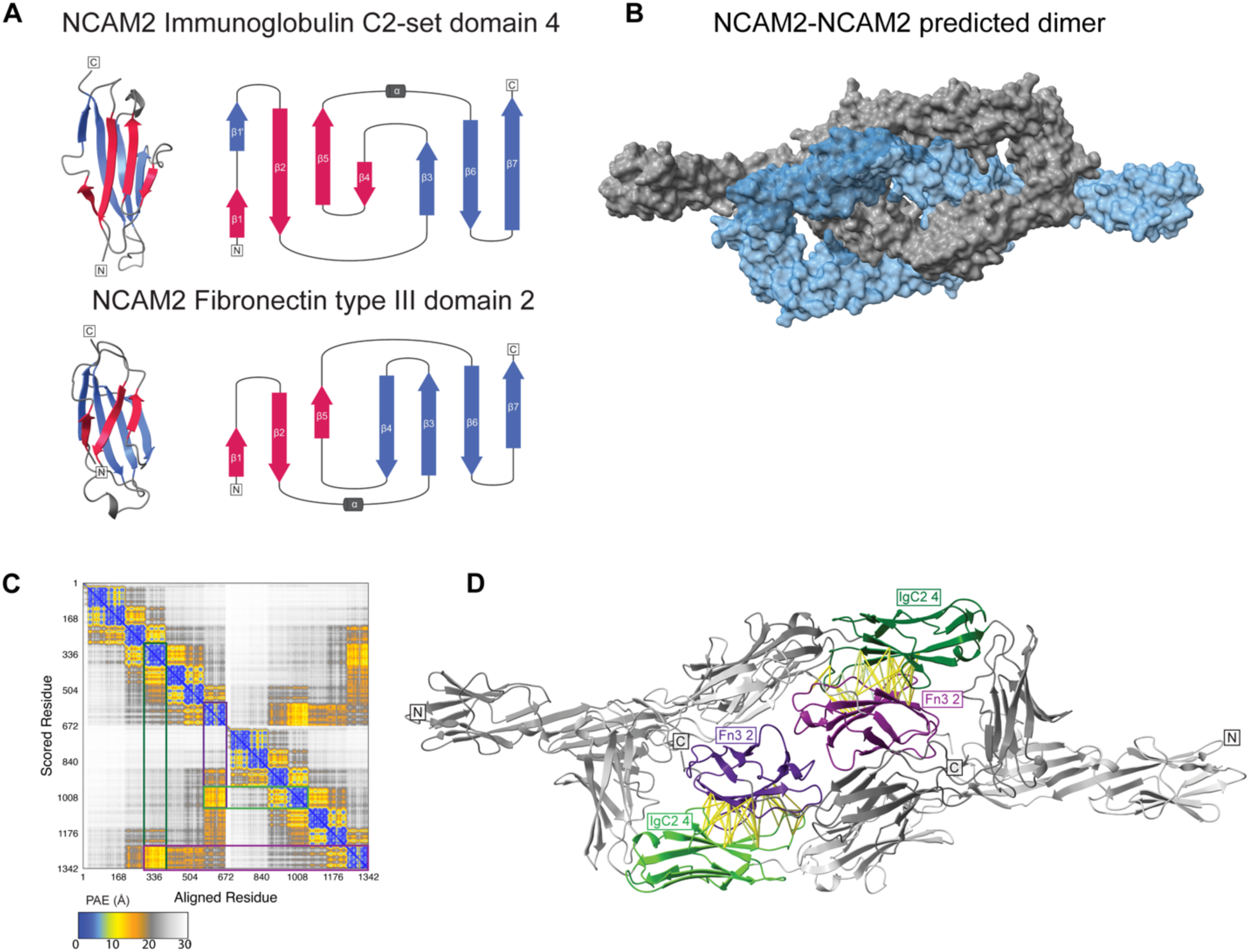
AlphaFold (v3) predictions of NCAM2 monomer and dimer structures. (A) Both immunoglobulin-like (IgC2) and fibronectin type III (Fn3) domains adopt the greek key motif composed of two β-sheets with antiparallel β-strands and an α-helix connecting the sheets (17). (B) Prediction of the NCAM2-NCAM2 dimer shows a tightly interlocking surface. Predicted TM-score (pTM) = 0.39, interface predicted TM-score (ipTM) = 0.39. (C) Predicted Aligned Error (PAE) matrix supports the dimer formation with the highest confidence interaction (between IgC2 set 4 and Fn3 domain 2) highlighted with bounding boxes. PAE provides a confidence measure for the position of any two amino acids within the structure. Blue-to-white gradient color block indicates high-to-low confidence. (D) Predicted NCAM2-NCAM2 dimer structure suggests interaction between IgC2 set 4 (colored green) and Fn3 domain 2 (colored purple) of opposing monomers. Predicted contacts are shown as rods between residues, filtered by interatomic distance ≤ 5 Å and PAE < 12 Å. Monomers are distinguished by lighter and darker shades.

